# Decoy Antibodies Block Extracellular HSP70, Prevent Self Signaling and Inhibit Melanoma Cell Survival

**DOI:** 10.64898/2025.12.29.696857

**Authors:** Noam Ben-Shalom, Shivang Parikh, Lilach Abramovitz, Ron Yefet, Paulee Manich, Dafna Tussia-Cohen, Ori Moskowitz, Roma Parikh, Stav Melamed, Ksenia Polonsky, Sebastien Apcher, Tzachi Hagai, Merav Cohen, Yochai Wolf, Ronnie Shapira-Frommer, Carmit Levy, Natalia T Freund

## Abstract

Melanoma cells actively secrete melanosomes-large, extracellular vesicles (EVs) enriched with oncogenic factors that reprogram the tumor microenvironment, enhance self-signaling, and promote tumor growth. Despite their abundance and immunogenic potential, humoral responses to melanoma-derived melanosomes remain unexplored. Here, we identify a novel immune surveillance mechanism in which melanosome-elicited decoy antibodies target melanoma-derived melanosomes by binding to the extracellular form of heat shock protein 70 (HSP70), a chaperone broadly implicated in cancer cell survival and stress adaptation. Anti-HSP70 decoy antibodies potently and effector-independently inhibit growth and survival of both murine and human melanoma cells and suppress key transcriptional programs involved in proliferation, cytoskeletal dynamics, and metabolism. In a preclinical B16 melanoma model, prophylactic administration of decoy monoclonal antibodies Mel322-34 and Mel321-35 conferred significant survival benefits of 27% and 48%, respectively. Strikingly, anti-HSP70 antibodies were enriched in the sera of melanoma patients achieving complete responses to immune checkpoint blockade, in contrast to non-responders with progressive disease. Collectively, our findings uncover a novel EV-antibody axis as a promising avenue to block cancer-promoting signaling pathways. Decoy autoantibodies targeting the extracellular form of HSP70 advance the understanding of tumor-intrinsic vulnerability and promote biomarker-driven immunotherapy in melanoma.

## INTRODUCTION

Melanoma is the most lethal of human skin cancers, accounting for the majority of skin cancer-related deaths, even though it comprises only a small percentage of all skin cancer cases [1–3]. Despite remarkable progress in therapy, including targeted treatments and immunotherapies, many patients develop resistance to these treatments over time [4–8]. Approximately 50% of patients with BRAF-mutated melanoma relapses within 1 year of starting combined BRAF/MEK inhibitor therapy [9]. Additionally, a significant proportion of patients fail to respond to immune checkpoint inhibitors or experience only transient benefits before disease progression [10]. Thus, a breakthrough in developing new strategies to prevent melanoma recurrence and metastasis is essential to overcoming treatment resistance and improving long-term outcomes for patients with advanced melanoma.

Immune response plays a crucial role in melanoma [11, 12]. Despite substantial evidence supporting T cells as key mediators of anti-melanoma immunity, either by directly killing tumor cells or activating other immune cells [13–15], the role of B cells in tumor response and surveillance remains less understood [16–18]. Studies suggest that tumor-infiltrating B cells are associated with improved survival [19–22] and enhanced responsiveness to immunotherapy [23, 24]. Furthermore, melanoma patients exhibit antibody responses, particularly autoantibodies against intracellular antigens such as tubulin, YWHAZ, MASP1, H4C1 and PF4, which are elevated in active disease [25]. Notably, Fässler M. *et al.* demonstrated that higher antibody titers against melanocyte differentiation antigens (TRP1, TRP2, gp100, and MelanA/MART1) correlated with improved responses to immune checkpoint inhibitors, prolonged progression-free intervals, and better overall survival [26]. Similarly, Stockert *et al.* reported that 9.4% of melanoma patients exhibited antibody responses against the cancer-testis antigen NY-ESO-1, a biomarker linked to improved survival [27]. However, these antibodies have primarily been investigated as diagnostic markers, and their functional role in melanoma immunity remains unclear.

Melanoma progression is characterized by the secretion of melanosomes by melanoma cells [28]. Melanosomes are specialized extracellular vesicles (EVs), 200–500 nm in size, produced by melanocytes and primarily responsible for melanin transport [29]. In cancer, melanosomes are reprogrammed to carry cancer-associated factors [30] and microRNAs that drive phenotypic changes in the tumor microenvironment, particularly in cancer-associated fibroblasts (CAFs) and stromal cells, thereby promoting tumor progression [28, 31–33]. Melanosomes are implicated in worsening disease outcomes [34] and contribute to drug resistance [35]. Although melanoma-derived melanosomes are large and enriched with cancer-associated proteins, they are restricted to the tissue, and their potential immunogenicity and the characteristics of antibodies targeting them have never been investigated.

In this study, we demonstrate for the first time that intravenous injection of melanoma-derived melanosomes to mice elicits protective B cell and antibody responses directed against the extracellular form of heat shock protein 70 (HSP70), a family of key regulators that promote self-signaling and proliferation of melanoma. Antibodies targeting this extracellular form of HSP70 inhibit melanoma cell growth, independently of additional immune effectors. Remarkably, we find that anti-HSP70 serum antibodies are naturally produced in melanoma patients, and those correlate with better response to immunotherapy. Our study unlocks the immunogenic potential of melanoma-derived melanosomes as a platform for melanoma vaccine, opening new promising avenues for therapeutic intervention.

## RESULTS

### Melanosome-Immunization Promotes Protection Against Melanoma in Mice

Melanosomes are secreted into the surrounding tissues *via* a paracrine mechanism [36]. To test whether melanoma melanosomes are immunogenic when administered systemically, we isolated melanosomes from B16 murine melanoma cell line [37] (see Materials and Methods) and immunized C57BL/6 mice (n=9) intravenously three times with melanosomes (Figure 1A). 19 days following Boost 2, we subcutaneously challenged the mice with 100,000 B16 melanoma cells. Mice immunized with melanosomes exhibited significantly prolonged survival compared to PBS control (p = 0.0019), with 33% of immunized mice surviving until end of the experiment (day 70), whereas none in the control group did (Figure 1B). Tumor onset in melanosome-immunized mice was delayed (average day 30 versus day 10 in controls), and tumors that did develop were smaller (Figure 1C-D). Serological analysis by ELISA revealed increasing production of melanosome-specific IgG (Figure 1E). Consistently, melanosome immunization induced a 3.6-fold increase in GC B cells (Figure 1F-G) and significant increases in CD4+ (1.5-fold) and CD8+ T cells (1.3-fold, Figure 1H-I), while natural killer (NK) cells remained unchanged (Supplementary Figure 1).

**Figure 1.**
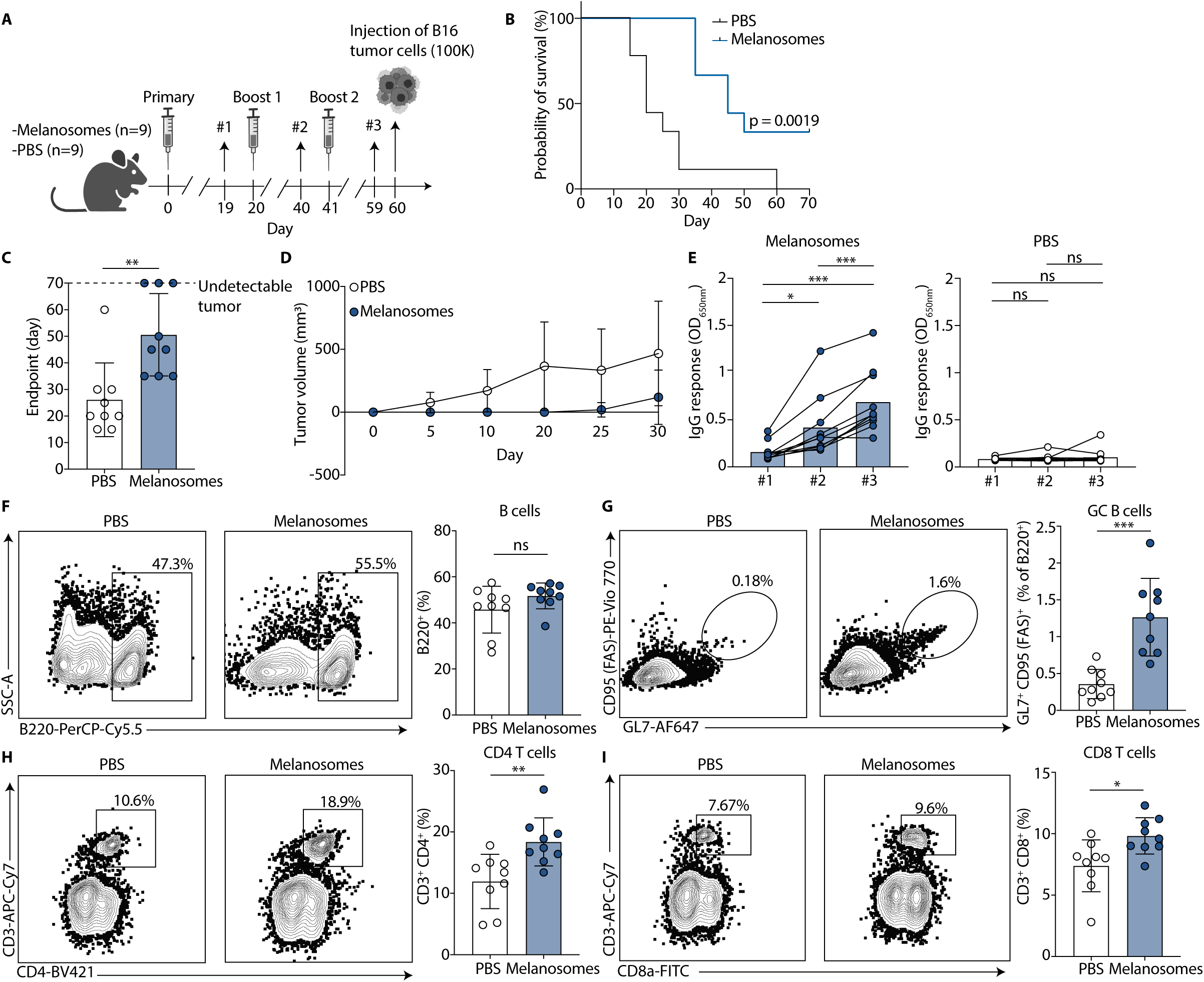
Melanosome-immunization promotes improved survival following melanoma-challenge. **A.** Schematic representation of the melanosome-immunization and melanoma challenge protocol. C57BL/6 mice were immunized with 1 µg melanosomes or PBS on day 0, followed by booster immunizations with 10 µg of melanosomes or PBS on days 20 and 41. On day 60, mice were challenged with a subcutaneous injection of 100,000 B16 melanoma cells. Blood samples were collected on days 19, 40, and 59 (n=9 per group). **B.** Kaplan-Meier survival curves comparing melanosome-immunized mice (blue, n=9) and PBS-immunized mice (black, n=9) after B16 melanoma challenge; the experiment was repeated 2 times. **C.** End point day for each mouse. Time to reach the tumor growth limit for individual mice in groups immunized melanosomes (n=9) and PBS (n=9). **D.** Tumor volume quantification (in mm³) over time in melanosome-immunized (blue circles, n=9) and PBS-immunized mice (white circles, n=9). Tumor volume was calculated as follows: V = 0.5 × Length × Width^2^. **E.** Serum IgG responses to melanosomes in melanosome-immunized mice (left, n=9) and in PBS-immunized controls (right, n=9); as measured by ELISA, presenting the raw O.D.650_nm_ at three time points: day 19 (#1), day 40 (#2), and day 59 (#3); the experiment was repeated 3 times. **F-I.** Flow cytometry plots of cell populations in the spleens of one representative melanosome-immunized and PBS-immunized mouse. The frequencies of the gated populations of all (n=9) mice in each group are presented in the right panels. The immune populations include- **F:** B cells (B220^+^), **G:** GC B cells (B220^+^ CD95 (FAS)^+^ GL7^+^), **H:** CD4^+^ T cells (CD3^+^ CD4^+^), and **I:** CD8^+^ T cells (CD3^+^ CD8^+^). The experiment was repeated 2 times. Graphs **C**, **D**, **F-I** present the meanL±Ls.d with each symbol represents an individual mouse. Graphs in **E** present the mean with each symbol represents an individual mouse. Statistical significance was determined using GraphPad Prism by Log-rank (Mantel-Cox) test in **B**, by Mann-Whitney U test in **C**, by Repeated measures one-way ANOVA with Tukey’s multiple comparisons test in **E**, and by unpaired two-sided Welch’s t-test in **F-I**. *, p < 0.05; **, p < 0.01; ***, *p* < 0.001, ****, p < 0.0001; “ns” indicates non-statistically significant differences.

### Anti-Melanosome mAbs Block Melanosome Signaling and Inhibit Growth

We investigated whether antibodies directed against melanosomes can mediate tumor suppressive functions and inhibit melanoma growth. Sera from melanosome-immunized mice (n=10) were collected on days 3, 5, 8, 10, 12 and 14 (Figure 2A). Anti-melanosome IgG became detectable by ELISA starting from day 8 post-immunization and continued to increase over time (Figure 2B). This binding was no longer detected when the melanosomes were trypsinized, indicating that the antibodies target membrane-associated proteins on the surface of melanosomes (Supplementary Figure 2). Next, we single cell sorted IgG^+^ GC B cells from spleens of four mice (Figure 2C-D), followed by IgH and IgK amplification and sequencing, as previously described [38, 39] (Supplementary Table 1). All mice exhibited expanded B cell clones, showing progressive accumulation of somatic hypermutations (SHM) and clonal bursts (Figure 2E and Supplementary Figure 3). We produced 13 monoclonal antibodies (mAbs), representing the different expanded B cell clonal families. Most mAbs exhibited binding to purified B16 melanosomes, as detected by ELISA and flow cytometry (Figure 2F-G). Among the mAbs tested, Mel321-31, Mel322-34, and Mel321-35 exhibited the strongest binding and were therefore selected for further studies.

**Figure 2.**
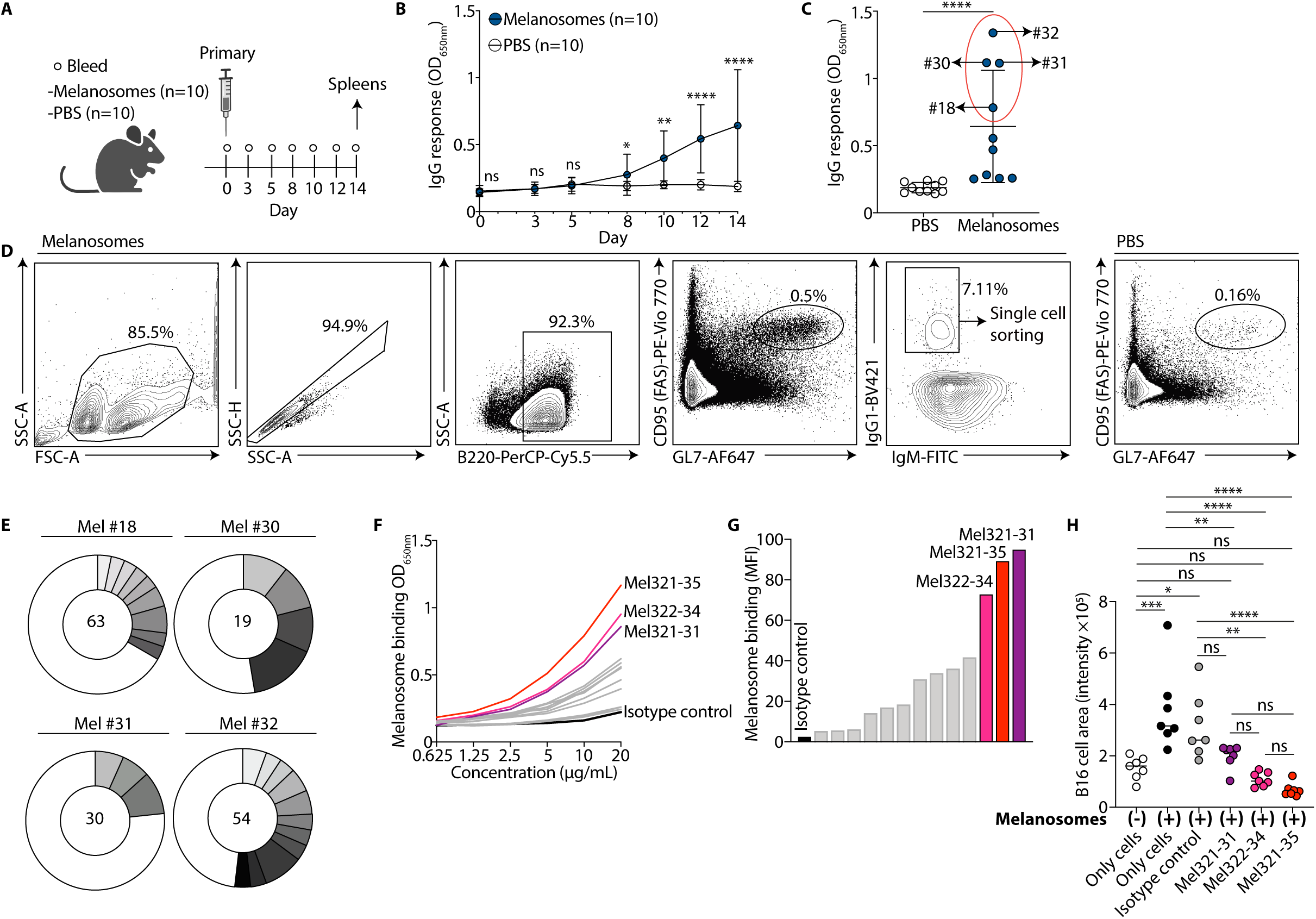
Melanosome-immunized mice show a profound B cell clonal expansion. **A.** Schematic representation of the melanosome-immunization protocol. C57BL/6 mice (n=10 per group) were immunized with 10 µg of melanosomes or PBS, and blood was collected on days 0, 3, 5, 8, 10, 12, and 14 following immunization. Spleens were harvested on day 14 post-immunization. **B.** Serum IgG response to melanosomes, measured by ELISA, in melanosome-immunized and PBS-immunized mice (n=10 per group). Group mean raw O.D. 650nm values at each time point (days 0, 3, 5, 8, 10, 12, 14); the experiment was repeated 3 times. **C.** Serum IgG response to melanosomes, measured by ELISA, in individual melanosome-immunized and PBS-immunized mice (n=10 per group) on day 14 post-immunization. The raw O.D. 650 nm values are presented. The four melanosome-immunized mice that were selected for B cell sorting (#18. #30, #31, and #32) are indicated by a red oval; the experiment was repeated 3 times. **D.** Left: Representative flow cytometry plot and gating strategy of IgG1^+^ GC B cells from a spleen of a melanosome-immunized mouse. The population that was single-cell sorted is indicated. Gating: lymphocytes → single cells → B cells (B220^+^) → GC B cells (CD95 (FAS)^+^ GL7^+^) → IgG1^+^ (IgG1^+^ IgM^−^). Right: GC B cells (CD95 (FAS)^+^ GL7+) staining from a spleen of a PBS-immunized mouse. **E.** Pie charts showing the number of paired IgG1^+^ Kappa sequences from the 4 melanosome-immunized mice (#18. #30, #31, and #32). The total number of sequences from each mouse is displayed in the center of each pie. Shaded slices indicate expanded B cell clones, while white slices represent unique sequences. **F.** Binding of mAbs to melanosomes (n = 13, 10 µg/mL), assessed by ELISA (the y axis shows raw O.D. 650 nm values). mGO.53.53 (black line) serves as an isotype control [58]. The three highest binding mAbs Mel321-31, Mel322-34, Mel321-35 are depicted in purple, magenta and red, respectively, while the other 10 mAbs are in gray. **G.** Binding of mAbs to melanosomes (2.5 µg/mL), as measured by flow cytometry (n=13 mAbs, 10 µg/mL) and detected by staining with anti-mouse IgG conjugated to Alexa Fluor 647. Median fluorescence intensity (MFI, y axis) was calculated for each mAb. The three highest binding mAbs Mel321-31, Mel322-34, Mel321-35 are depicted in purple, magenta and red, respectively, while the other 10 mAbs are in gray. mGO.53 is in black and serves as an isotype control; the experiment was repeated 3 times. **H.** Quantification of fluorescence intensity of TdT-expressing B16 cells under different treatment conditions: untreated cells (no melanosomes, white circles), cells with 5 µg/mL of external melanosomes (black circles), cells treated with isotype control, mGO.53 (25 µg/mL) plus 5 µg/mL of external melanosomes (gray circles), Mel321-31 (25 µg/mL) plus 5 µg/mL of external melanosomes (purple circles), Mel322-34 (25 µg/mL) plus 5 µg/mL of external melanosomes (pink circles), and Mel321-35 (25 µg/mL) plus 5 µg/mL of external melanosomes (red circles). n=7 per treatment group; the experiment was repeated 3 times. Graphs **B-C** present the meanL±Ls.d with each symbol represents an individual mouse. Statistical significance was determined using GraphPad Prism by Mann-Whitney U test in **B-C** and by one-way analysis of variance (ANOVA) with Tukey’s multiple comparisons post-test in **H**. *, p < 0.05; **, p < 0.01; ***, *p* < 0.001; ****, p < 0.0001; “ns” indicates non-statistically significant differences.

Purified B16 melanosomes had a striking effect on B16 melanoma cells, markedly increasing fluorescence intensity by promoting greater cell adherence. Within just 2 hours of culture, cells exposed to melanosomes exhibited a 148% increase in intensity compared to untreated controls (Figure 2H, black vs. empty circles). While melanosomes have previously been shown to support the growth of fibroblasts and tumor-associated stromal cells, our findings reveal that they also enhance the proliferation of the melanoma cells that secrete them. This autocrine-like ‘melanosomal effect’ was abolished when melanosomes were pre-incubated with monoclonal antibodies Mel322-34 or Mel321-35 (Figure 2H, Supplementary Figure 4). Partial inhibition was observed with Mel321-31. These results indicate that melanosome-targeting antibodies are acting as decoys that block the melanosomal growth-promoting effects.

### Anti-melanosome mAbs inhibit B16 melanoma cell survival

MAbs Mel321-31, Mel322-34, and Mel321-35 significantly impaired the growth of TdT-expressing B16 melanoma cells over 60 hours, reducing cell growth by 4.1-fold, 8.4-fold, and 8.2-fold, respectively, relative to the isotype control, as quantified by B16 fluorescence intensity (Figure 3A, Supplementary Figure 5). Incubation of B16 cells with melanosome-targeting mAbs increased caspase-3/7 activity detected using an activated fluorescent DNA dye, suggesting that the mAbs induced apoptosis (Figure 3B). To further explore this, we performed mRNA-seq analysis. We identified 23 significantly downregulated genes shared among B16 cells treated with Mel321-31, Mel322-34, and Mel321-35, versus cells cultured with isotype control. Amongst these, 12 were related to apoptosis and survival, 5 were related to cytoskeleton organization, 3 to metabolism and 3 to stress response (See Materials and Methods, Figure 3C). Gene set enrichment analysis (GSEA) identified only significantly downregulated pathways in mAbs-treated cells. Notably, the IL6/JAK/STAT3 signaling pathway was consistently and significantly suppressed across all three mAbs (Figure 3D). This pathway plays a critical role in tumor cell proliferation, survival, invasion, and metastasis [40]. Reduction in IL-6 following mAb treatment was confirmed by ELISA, although for Mel321-31 the reduction in IL-6 did not reach statistical significance (Figure 3E). These results support the observation that Mel321-31, Mel322-34, and Mel321-35 change the cellular transcriptional landscape.

**Figure 3.**
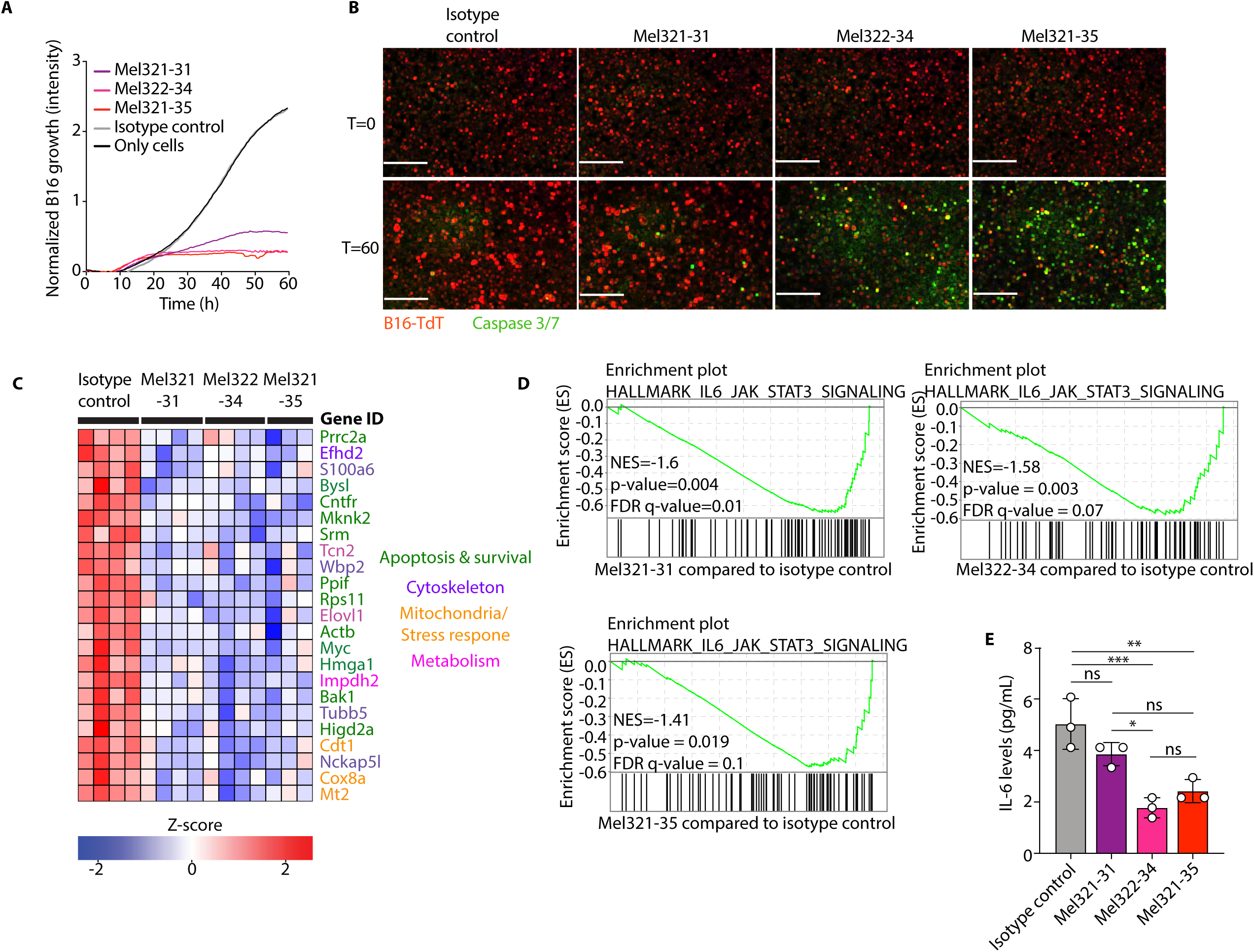
Anti-melanosome mAbs inhibit melanoma cell growth. **A.** Quantification intensity of 15 × 10^3^ TdT-expressing B16 cells over 60 hours following treatment with mAbs (25 µg/mL, 6 repeats for each mAb), measured by IncuCyte. No mAb (‘Only cells’, black) and mGO.53 (isotype control, gray) serve as controls. The three mAbs Mel321-31, Mel322-34, Mel321-35 are depicted in purple, magenta and red, respectively. The experiment was repeated 3 times. **B.** Representative images of TdT-expressing B16 cells (red) merged with Caspase 3/7 activated cells (green) at 0 and 60 hours post-treatment. Scale bars: 400 µm. **C.** Heatmap showing z-scaled log-normalized counts per million for shared downregulated differentially expressed genes in B16 cells after 24-hour treatment with mAbs Mel321-31 (n=4), Mel322-34 (n=4), Mel321-35 (n=3), and isotype control, mGO.53 (n=4). **D.** GSEA of downregulated pathways shared among B16 cells following 24-hour treatment with mAbs Mel321-31, Mel322-34, and Mel321-35 compared to isotype control, mGO.53. **E.** IL-6 quantification by ELISA. mAbs (25 µg/mL) - Mel321-31 (n=3), Mel322-34 (n=3), Mel321-35 (n=3), and mGO.53 (n=3) - were incubated with B16 cells. The supernatant was collected after 40 hours and analyzed for IL-6 levels; n=3, the experiment was repeated 2 times. Graph **E** presents the mean ±Ls.d with each symbol represents an individual replicate. Statistical significance was determined using GraphPad Prism by one-way analysis of variance (ANOVA) with Tukey’s multiple comparisons post-test in **E**. *, p < 0.05; **, p < 0.01; ***, *p* < 0.001; “ns” indicates non-statistically significant differences.

### Anti-Melanosome mAbs Prolong Survival in B16-Challenged Mice

To estimate the ability of the mAbs to inhibit tumor development *in vivo*, we administered 75 µg of antibody (Mel321-31, n=10, Mel322-34, n=10, Mel321-35, n=10) or the isotype control (n=10) intraperitoneally to C57BL/6 mice 1 hour before injecting subcutaneously 100,000 B16 melanoma cells. A second dose of 35 µg mAb was administered one week after tumor injection, and the mice were monitored for tumor growth and survival (Figure 4A). Mice treated with Mel322-34 and Mel321-35 showed a significant increase in survival (27 % increase and 48 % increase, respectively), compared to mice treated with isotype control. Mice treated with Mel321-31 had similar survival rates to mice treated with isotype control (Figure 4B). Notably, treatment with Mel321-35 led also to a significant delay in tumor onset in the mice (average of 15.5 days in isotype control compared to average of 22.9 days in Mel321-35, Figure 4C). The mAbs reduced tumor volume and inhibited tumor growth compared to mice treated with isotype control (Figure 4D-E). Focusing on Mel321-35, which conferred the strongest survival benefit, we found no major differences in splenic macrophage, NKLcell, or CD4L and CD8L T-cell frequencies (Supplementary Figure 6A-D). In contrast, the melanosome-targeting mAb reduced splenic B-cell frequencies (Figure 4F). This reduction was accompanied by an increase in activated B cells (B220L CD69L) within the tumor-draining lymph node (TDLN) relative to controls, without changes in overall tumor immune infiltration (CD45L) (Figure 4G and Supplementary Figure 6E-F). Together, these findings suggest that treatment selectively enhances B-cell migration to, and activation within, the tumor-proximal lymphoid niche.

**Figure 4.**
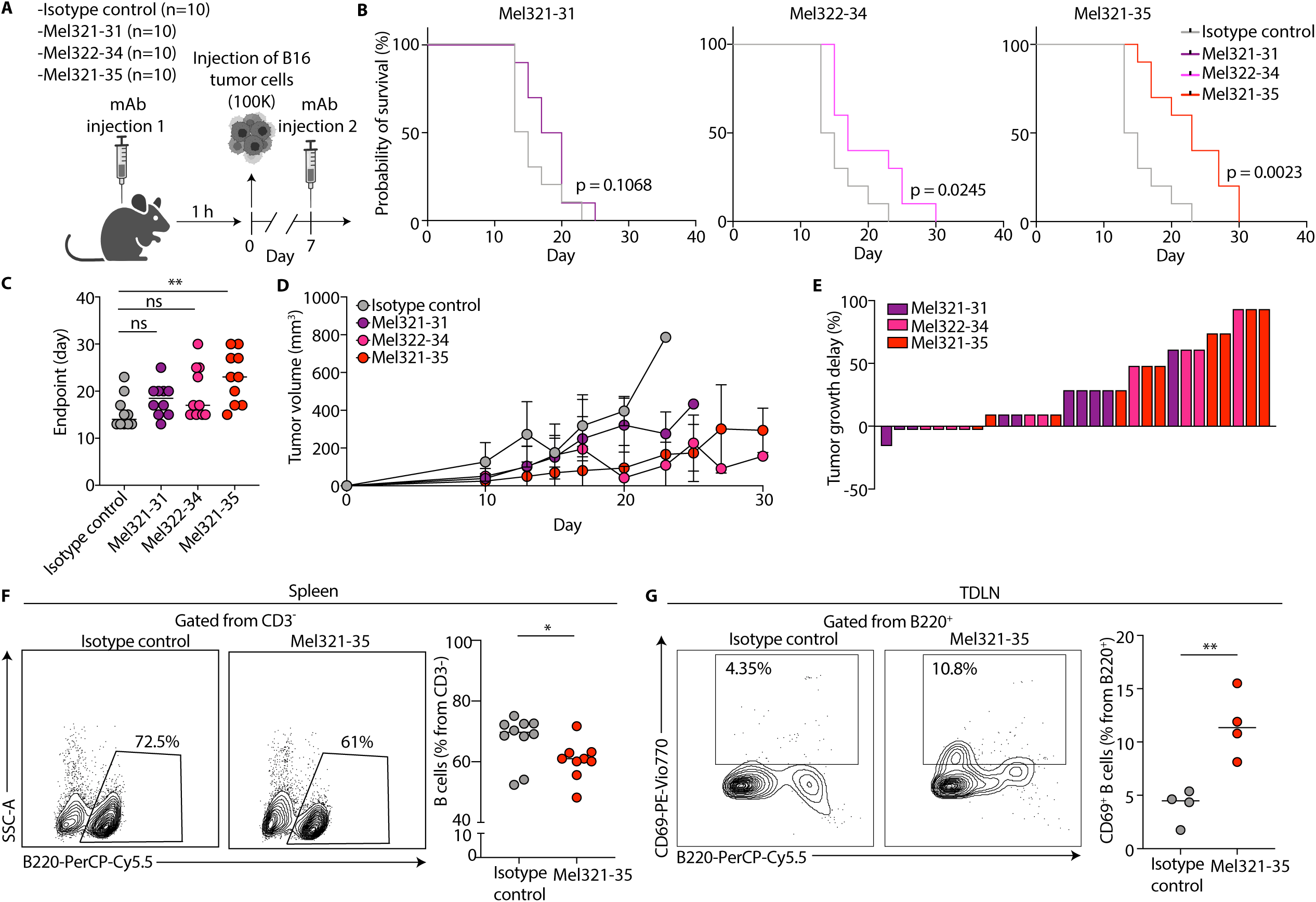
Anti-melanosome mAbs prolong survival in B16 challenged mice. **A.** Schematic representation of the passive immunization and tumor challenge strategy. Mice were immunized with 70 µg of the following mAbs: Mel321-31, Mel322-34, Mel321-35, and isotype control, mGO.53. After one hour, 100,000 B16 cells were injected subcutaneously into all mice. One week following the mAb and B16 injection, another boost of 35 µg of each mAb was administered to each mouse (n=10 per group). **B.** Kaplan-Meier survival curves comparing mice treated with mAbs Mel321-31 (purple line, n=10), Mel322-34 (pink line, n=10), and Mel321-35 (red line, n=10) to the isotype control, mGO.53 (gray line, n=10). **C.** End point day for each mouse. Time to reach the tumor growth limit for individual mice in groups immunized with Mel321-31 (purple circles, n=10), Mel322-34 (pink circles, n=10), Mel321-35 (red circles, n=10), and isotype control, mGO.53 (gray circles, n=10). **D.** Tumor volume quantification (in mm³) over time in mice treated with Mel321-31, Mel322-34, Mel321-35, and isotype control, mGO.53 (n=10 per group). Tumor volume was calculated as follows: V = 0.5 × Length × Width^2^. Color code is the same as in **C**. **E.** Tumor growth delay (TGD) frequencies for individual mice in the Mel321-31 (purple bars, n=10), Mel322-34 (pink bars, n=10), and Mel321-35 (red bars, n=10) compared to the average of the group of isotype control, mGO.53-immunized mice (n=10). Calculated as TGD% = [(T_Treated_ − T_Control_) / T_Control_] × 100, where T_Control_ is the mean endpoint of the mGO.53-immunized group, and T_Treated_ is the endpoint for each mouse in the Mel321-31, Mel322-34 and Mel321-35 immunized groups. **F.** B cell frequency in spleens of treated mice. The left panel shows representative flow cytometry plots of B220^+^ splenocytes of one Mel321-35- and isotype control- treated mouse (gated from CD3^−^, Supplementary Figure 6), while the right panel shows quantification of the gated population (Mel321-35, n=8, isotype control, n=9). **G.** Activated B cell frequency in TDLN of treated mice. The left panel shows representative flow cytometry plots of B220^+^ CD69^+^ (gated from B220^+^, Supplementary Figure 6) of one Mel321-35- and isotype control- treated mouse, while the right panel shows quantification of the gated population (Mel321-35, n=4, isotype control, n=4). Graphs **C, F, G** present the mean and graph **D** presents the mean ±Ls.d with each symbol represents an individual mouse. Statistical significance was determined using GraphPad Prism by Log-rank (Mantel-Cox) test in **B**, by one-way analysis of variance (ANOVA) with Tukey’s multiple comparisons post-test in **C** and by unpaired two-sided Welch’s t-test in **F-G**. *, p < 0.05; **, p < 0.01; “ns” indicates non-statistically significant differences.

### Identification of HSP70 as the Target of Anti-Melanosome mAbs

Western blot analysis of melanosome lysates-containing both surface and internal proteins-revealed that all three mAbs recognized a protein of approximately 70 kDa (Figure 5A). To identify the target, immunoprecipitated proteins were analyzed by liquid chromatography-tandem mass spectrometry (LC-MS/MS), which revealed four highly enriched members of the HSP70 family: HspA1A, HspA1L, HspA5, and HspA8 (Figure 5B-C, Supplementary Figure 7). ELISA and SPR analyses confirmed that Mel321-31, Mel322-34, and Mel321-35 bound to HSP70 family members with measurable affinities (9.87×10^−9^, 5.52×10^−9^, 3.1×10^−9^, Figure 5D and Supplementary Figure 8). HSP70 proteins are well-known regulators of apoptosis and cell survival and are frequently overexpressed in melanoma and other cancers [41, 42]. HSP70 is a potent inhibitor of apoptosis, a known activator of the JAK/STAT pathway, and a key suppressor of caspase-3 activity [43–45]. These functional roles align with our earlier findings, in which antibody treatment inhibited cell survival and suppressed IL-6 secretion—further supporting HSP70 as the principal target of the mAbs.

**Figure 5.**
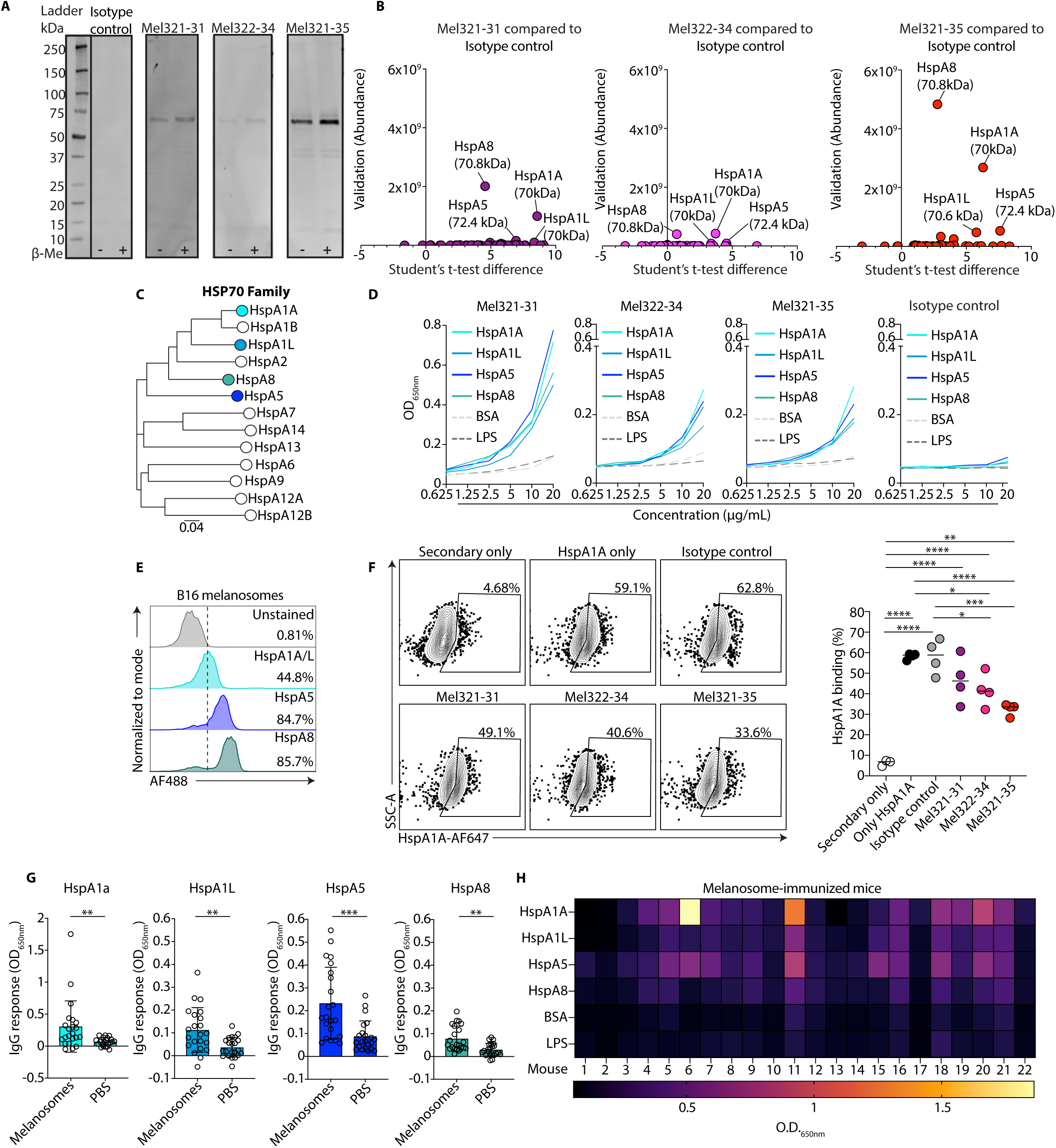
Identification of HSP70 as the target of anti-melanosome mAbs. **A.** Western blot analysis showing the binding of mAbs Mel321-31, Mel322-34, Mel321-35, and the isotype control, mGO.53 to a whole lysate of melanosome proteins. 15 µg of whole lysate was loaded on a protein gel with or without β-Mercaptoethanol, followed by incubation with 25 µg/mL of each mAb (Mel321-31, Mel322-34, Mel321-35, or isotype control, mGO.53) and 1:5000 anti-mouse secondary antibody; the experiment was repeated 3 times. **B.** Mass spectrometry analysis showing enriched proteins following immunoprecipitation of proteins from the whole lysate of melanosomes using mAbs Mel321-321 (purple circles, n=4), Mel322-34 (pink circles, n=4) and Mel321-35 (red circles, n=4) compared to isotype control, mGO.53 (n=4). HspA1A, HspA1L, HspA5, HspA8 and their molecular weight are indicated; the experiment was repeated 2 times. **C.** Phylogenetic tree showing the mouse HSP70 protein family. The bar at the bottom of the tree provides a scale. **D.** Binding of mAbs Mel321-31, Mel322-34, Mel321-35, and isotype control, mGO.53 to HspA1A, HspA1L, HspA5, HspA8, and two control proteins (BSA and LPS), as measured by ELISA. O.D. values at 650 nm are shown for six antibody dilutions (20, 10, 5, 2.5, 1.25, 0.625 µg/mL); the experiment was repeated 4 times. **E.** Histograms showing the expression of HspA1A/L, HspA5, and HspA8 on B16 murine melanosomes, assessed by flow cytometry. The dashed line indicates the threshold for positive staining. **F.** HspA1A binding to B16 cells. Left: Representative flow cytometry plots showing the binding frequency of AF647-conjugated HspA1A to B16 cells under different treatment conditions: Secondary only control (Streptavidin-AF647, n=3), HspA1A alone (n=4), and pre-incubation of AF647-conjugated HspA1A with mAbs- isotype control (n=4), Mel321-31 (n=4), Mel322-34 (n=4), and Mel321-35 (n=4). Right: Quantification of binding frequencies; the experiment was repeated 2 times. **G.** Reactivity of IgG antibodies in sera from melanosome-immunized (n=22) and PBS-immunized (n=22) mice as measured by ELISA against HspA1A, HspA1L, HspA5, and HspA8. Raw O.D.650 nm values were obtained after subtracting background binding to BSA; Every data point is one mouse, the experiment was repeated 2 times. **H.** Heat map representing the values in **G** for each mouse along with reactivity to BSA and LPS. Graph **F** presents the mean with each symbol represents an individual replicate and the graphs in **G** present the mean ±Ls.d with each symbol represents an individual mouse. Statistical significance was determined using GraphPad Prism by one-way analysis of variance (ANOVA) with Tukey’s multiple comparisons post-test in **F**, and by Mann-Whitney U test in **G**. *, p < 0.05; **, p < 0.01; ***, *p* < 0.001, ****, p < 0.0001; “ns” indicates non-statistically significant differences.

Flow cytometry using commercial antibodies confirmed high surface expression of HspA1A/L (44.8%), HspA5 (84.7%), and HspA8 (85.7%) on B16 murine melanosomes (Figure 5E). Mel322-34 and Mel321-35 significantly inhibited the binding of fluorescently labeled HspA1A to B16 cells, while Mel321-31 showed a non-significant effect (Figure 5F). AlphaFold3 modeling predicted (exhibiting high-confidence, predicted TM-score > 0.8) that both mAbs Mel321-31 and Mel322-34 majorly target the nucleotide-binding domain (NBD) across HspA1A, HspA1L, HspA5, and HspA8. Mel321-35, which consistently demonstrated stronger functional activity, exhibited broader specificity, interacting with both the NBD and the substrate-binding domain (SBD) of HspA1A, HspA1L, HspA5 and HspA8 (Supplementary Figure 9). Finally, we tested the reactivity of sera derived from melanosome-immunized mice against HSP70 family members. As expected, the sera of immunized mice reacted against all four HSP70 family members, in contrast to the sera from PBS-immunized mice, which showed no such reactivity (Figure 5G-H). These results indicate that Mel321-31, Mel322-34, and Mel321-35 function as decoy antibodies, binding melanosome-associated HSP70 and blocking its interaction with melanoma cells, thereby disrupting pro-survival signaling.

### Anti-HSP70 Antibodies Correlate with Better Response to Therapy

To assess the relevance of our mAbs to targeting human melanoma, we compared the expression of HSP70 family members in MNT-1 melanoma cell line. All four HSP70 family members were increased in their expression relative to healthy fibroblasts (Figure 6A). HSP70 expression was further confirmed by flow cytometry of purified MNT-1 melanosomes demonstrating high levels of HspA1A/L (67.6%), HspA5 (64.5%), and HspA8 (66.3%) on the surface of MNT-1 melanosomes (Figure 6B). We next examined whether our mAbs block the survival of MNT-1 cells similarly to what we observed in murine B16 cells. All three mAbs, Mel321-31, Mel322-34, and Mel321-35, induced apoptosis in MNT-1 (Figure 6C), compared to primary human fibroblasts where no cell death was detected (Figure 6D). Taken together, these results align with prior reports showing strong cross-species conservation of HSP70 family members [46–48], which likely enables shared recognition by our mAbs.

**Figure 6.**
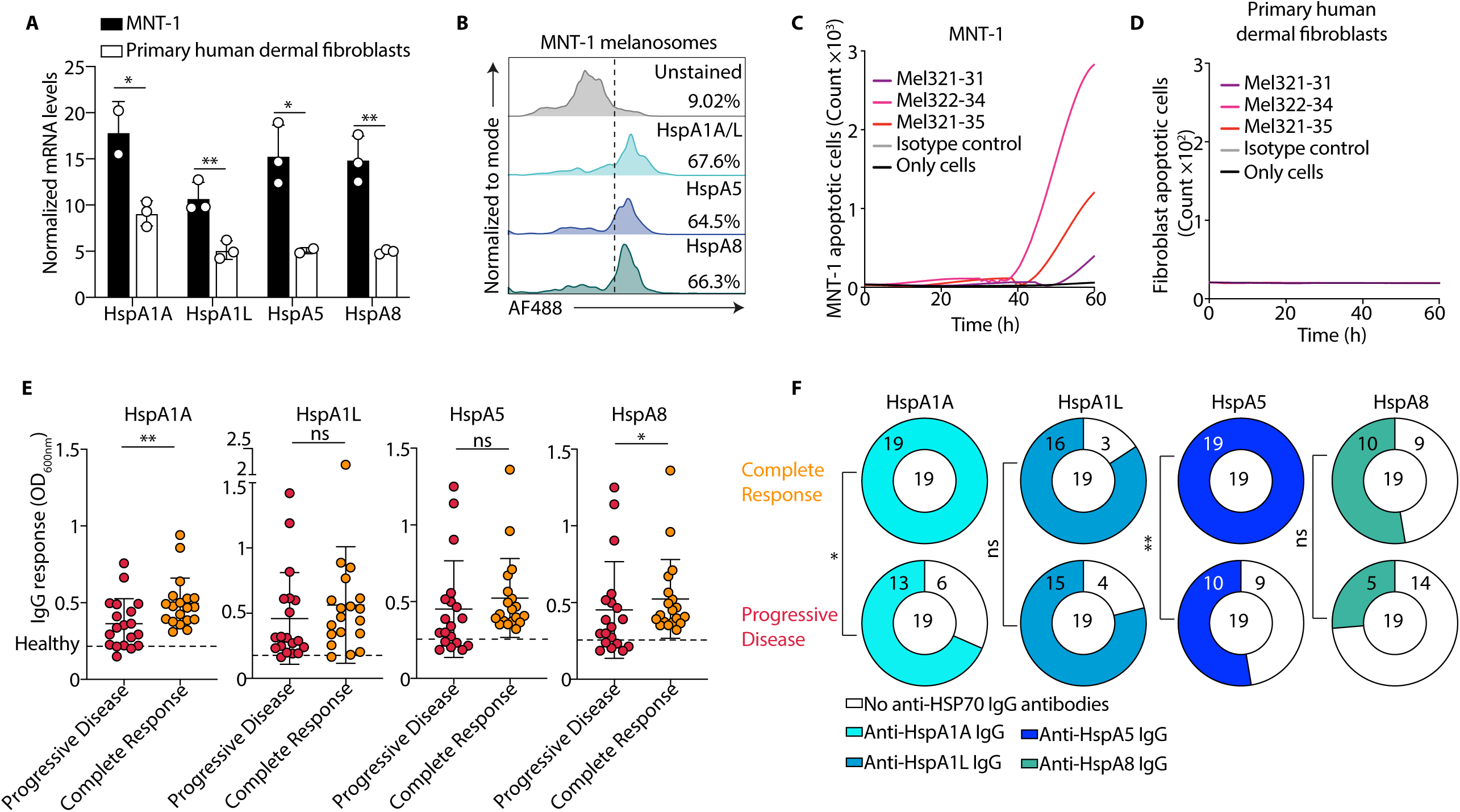
Anti-HSP70 serum response correlates with better response to therapy. **A**. Levels of HSP70 family members (HspA1A, HspA1L, HspA5 and HspA8) mRNA in MNT-1 and primary human dermal fibroblasts, as measured by real-time PCR (n=2-3). The results were normalized against the housekeeping gene RPLP0. **B.** Histograms showing the expression of HspA1A/L, HspA5, and HspA8 on MNT-1 human melanosomes, assessed by flow cytometry. The dashed line indicates the threshold for positive staining. **C-D:** Quantification of cell death in MNT-1 cells (**C**) as well as in primary human dermal fibroblasts (**D**) over 60 hours following treatment with mAbs (25 µg/mL, 3 repeats for each mAb), measured by IncuCyte. No mAb is in black, mGO.53 is an isotype control and is shown as a gray line. The three mAbs Mel321-31, Mel322-34, Mel321-35 are depicted in purple, magenta and red, respectively. mAbs were incubated with 15 × 10^3^ cells, followed by the addition of a caspase 3/7 dye to detect apoptosis; the experiment was repeated 3 times. **E.** Reactivity of IgG antibodies in sera from patients in the “Progressive Disease” group (red circles, n=19) and the “Complete Response” group (orange circles, n=19), assessed by ELISA against HspA1a, HspA1L, HspA5, and HspA8. Raw O.D.600 nm values were obtained after subtracting background binding to BSA. The dashed line represents the mean response of sera from healthy donors (n=8). **F.** Pie charts illustrating the proportion of patients who developed an IgG response against HspA1a, HspA1L, HspA5, or HspA8. The colored portion represents patients with a detectable IgG response, while the white portion represents those without a response. The top row represents the Complete Response group (n=19), and the bottom row represents the Progressive Disease group (n=19). The threshold for a positive response was defined as greater than the mean response of healthy donors (n=8) + 0.05 O.D.600nm. Graphs A presents the mean ±Ls.d and graph **E** presents the mean ±Ls.d with each symbol represents an individual melanoma patient. Statistical significance was determined by unpaired two-tailed Student’s t-test **A**, Mann-Whitney U test in **E**, and Fisher’s exact test in **F**. *, p < 0.05; **, p < 0.01; “ns” indicates non-statistically significant differences.

Lastly, we investigated whether melanoma patients develop serum antibody responses to HSP70. Of the 38 stage 3 and 4 inoperable metastatic melanoma patients treated with immune checkpoint inhibitors, 19 achieved a complete response with tumor regression (“Complete Response”), while the remaining 19 showed no response, with continued disease progression (“Progressive Disease”). The Complete Response group exhibited significantly stronger IgG responses to HspA1A and HspA8 compared to the Progressive Disease group (40% and 18.7% increase, respectively, Figure 6E). Furthermore, 100% of the patients in the Complete Response group developed antibodies to HspA1A and HspA5 compared to Progressive Disease group where only 68% of the patients developed antibodies to HspA1A, and 52% to HspA5 (Figure 6F). These findings suggest a potential association between the presence of anti-HSP70 antibodies and responsiveness to immune checkpoint therapy, indicating a possible functional role for these antibodies in controlling melanoma.

## DISCUSSION

Melanoma remains a clinical challenge due to its high metastatic potential, adaptability to diverse microenvironments, and resistance to current therapies [3, 49, 50]. Recent insights reveal that melanosomes—normally involved in pigment transfer—are repurposed by melanoma cells into potent cancer-promoting agents. Much like viruses, these extracellular vesicles disseminate pro-tumorigenic signals to surrounding cells and reinforce survival pathways, contributing to cancer spread. However, their autocrine role in supporting melanoma cells was not demonstrated. Our study reveals that melanoma-derived melanosomes directly enhance melanoma cell growth, acting as autonomous pro-survival units. Targeted blockade using melanosome decoy antibodies completely abrogated this effect. Remarkably, the 5-day antibody treatment not only disrupted melanosome-driven growth and invasion but also induced apoptotic responses. These results uncover a critical self-sustaining mechanism in melanoma—a new vulnerability that can be effectively targeted by antibodies.

While melanosomes and melanogenesis have long been associated with tumor progression and metastasis [32, 33, 51–53], their capacity to provoke an adaptive immune response has remained unexplored. Unlike other EVs, melanosomes are typically confined to the tumor microenvironment [28, 31]. Here, we show that systemic immunization with melanoma-derived melanosomes elicits a robust B and T cell response, leading to prolonged survival and reduced tumor burden in mice. These findings shift the conventional understanding of melanosomes beyond their classical function in pigment transport [29, 54], and pathological function in immune niche deprogramming [28], positioning them as immunogenic entities and targets for antibodies with direct implications for melanoma therapy.

A key outcome of our study is the identification of HSP70 proteins as central targets of the anti-melanosome antibody response. While previous work—such as that by Komarova *et al*.—has shown that immunization with HSP70-enriched EVs can reduce tumor growth and improve survival in a B16 melanoma model, those effects were NK cell-mediated, and the elicited antibodies lacked intrinsic activity [55]. In contrast, our study reveals that melanosome-targeting antibodies induce apoptosis independently of NK cells. Indeed, we observed no changes in NK cell frequencies following melanosome immunization or after passive transfer of anti-melanosome mAbs. Notably, the antibodies act directly upon binding—akin to neutralizing antibodies targeting viral antigens—underscoring a unique autonomous mechanism of action. Another study reports anti-HSP70 mAb isolation by injecting 14-mer HSP70-derived peptides into mice and generating hybridomas [56]. Similar to what we describe, the generated mAb cmHsp70.1 induced anti-melanoma activity. However, here too, the activity of cmHsp70.1 relied on effector cell-mediated killing mechanisms [56]. In contrast, we are the first to report that anti-HSP70 mAbs elicited following melanosome immunization mediate melanoma cell killing without any additional effector cells.

HSP70 is highly conserved across a broad range of species, which likely explains the observed cross-reactivity of the antibodies between murine and human melanoma cell lines [46–48]. Furthermore, high HSP70 expression has long been associated with poor prognosis across multiple cancer types, underscoring its clinical relevance in tumor progression [57]. Yet, despite extensive research on its role in oncogenesis, the presence and significance of autoantibodies against HSP70 in cancer patients have remained largely unexplored. In this study, we uncover a robust autoantibody response to HSP70 in melanoma patients, which strongly correlates with favorable clinical outcomes and complete response to immune checkpoint blockade therapy. Interestingly, we observed elevated anti-HSP70 antibody levels in complete responders. Notably, anti-melanosome antibody levels did not differ between responders and non-responders (Supplementary Figure 10), suggesting that the anti-HSP70 response is not a result of general autoreactivity. Instead, these findings raise the possibility that anti-HSP70 antibodies contribute functionally to antitumor immunity, potentially through synergy with other immune cells, as well as directly impairing tumor cell survival. Deeper mechanistic studies investigating how ICI-enhanced CD8L T-cell activity synergizes with antibodies targeting HSP70 are required to determine whether HSP70-targeting antibodies have predictive or therapeutic relevance in melanoma and other HSP70-expressing cancers.

## Materials and Methods

### Ethics statement

#### Human samples

38 Participants were recruited from Sheba Medical Center under clinical study protocol SMC-2437-15, following approval from the Institutional Review Board (IRB) and adherence to the Helsinki Declaration guidelines. Patient selection was based on clinical diagnosis, and all participants provided written informed consent before inclusion in the study. The study was approved by the Institutional Ethics Committee of Tel Aviv University under protocol number 0009876-1. Additionally, a cohort of healthy donors, with no history of melanoma, was recruited as a control group for baseline antibody reactivity comparisons. Experiments with sera from healthy donors obtained from the Israeli Blood Bank (approval number: 0002682-1). The criteria for healthy donors include screening for blood type and for antibodies against the following infectious diseases: Syphilis (TPHA) HIV (HIV I/II) Hepatitis C, Hepatitis B, Human T-lymphotropic virus, and West Nile virus.

#### Animal studies

All housing conditions and all experimental procedures were approved and monitored by the Institutional Animal Care and Use Committee (IACUC) of Tel Aviv University. IACUC permit: 01-15-086 and 01-19-003. Animal experiments were conducted using female C57BL/6J mice (8–12 weeks old; age-matched within experiments) obtained from ENVIGO Laboratories (Jerusalem, Israel). Mice were housed in groups of 3–4 per cage under standard vivarium conditions (22 ± 2 °C, 12-hour light/dark cycle) with ad libitum access to food and water. Experimental groups were matched for age, weight, and drug administration. To minimize procedural stress, animals were acclimated to the vivarium for at least 3 weeks prior to experimentation.

#### Mice immunization and tumor model

C57BL/6 mice were injected with 1 or 10 µg of freshly purified melanosomes (see below “Melanosome Purification”) *via* intravenous injection. To induce tumor formation, 100,000 B16 melanoma cells were injected subcutaneously into each mouse. Tumors were measured weekly using digital caliper (Sangabery) and tumor volume was calculated using the formula: V = 0.5 × l × W ^2^, where V represents tumor volume, l is tumor length, and W is tumor width. For treatment with mAbs, 70 µg of each mAb (Mel321-31, Mel322-34, Mel321-35, and isotype control, mGO.53) was injected one hour prior to tumor cell injection. An additional boost of 35 µg of each mAb was administered one week after the initial tumor challenge. Mice were sacrificed if the tumor diameter reached ≥15 mm (absolute limit) in the melanosome-immunization model, ≥10 mm (absolute limit) in the prophylactic model, or if any of the following occurred: ulceration, necrosis, infection, severe weight loss (>20%), or signs of severe distress or impaired mobility.

#### Extraction of blood, spleens, tumor draining lymph nodes and tumors

For small blood samples (up to 50 µL) used in serological analyses, blood was collected from the cheek vein of live animals. For whole blood collection, animals were euthanized, and approximately 1 mL of blood was withdrawn directly from the heart. The blood was incubated at room temperature for 30 minutes to promote clotting, then centrifuged at 2000 g for 10 minutes at 4 °C to separate the serum. The serum was collected and stored at −20 °C for further analysis. Spleens and tumor draining lymph nodes were harvested and washed with 1X PBS. The tissues were then mechanically disrupted using a pestle (Biofil), passed through a 70 µm cell strainer (Corning), and centrifuged at 400 g for 5 minutes at 4 °C. For spleens, red blood cells were lysed by incubating the cell suspension in red blood cell lysis buffer (155 mM NH4Cl, 10 mM KHCO3) for 5 minutes at room temperature. After lysis, the cells were washed with 1X PBS and centrifuged again at 400 g for 5 minutes at 4 °C. The supernatant was discarded, and the cells were resuspended in fetal bovine serum (FBS; Thermo Fisher Scientific) with 10% DMSO (Sigma) before being frozen at −80 °C for further analysis. Tumors were processed using multi tissue dissociation kit 1 according to the manufacturer’s instructions (Milteny Biotec, 130-110-201).

#### Melanosome purification

Melanosomes were isolated from B16 cells as previously described [28, 33]. Briefly, cells are grown to 60% confluence, media was changed, and they were grown for 5 days. After 5 days, the conditioned media was collected and centrifuged at 300xg for 15 minutes followed by 1000xg for 30 minutes. Melanosomes were then pelleted by centrifuging the supernatant at 20,000xg for 70 minutes. Melanosomes were resuspended in PBS and used as described further. Melanosome trypsinization was made as previously [33]. The melanosome isolation protocol was calibrated and validated according to MISEV2023 guidelines. Melanosome and exosome size distributions were determined using NanoSight nanoparticle tracking analysis, with melanosomes defined within the 200–500 nm range, and exosomes within 70–100 nm. Differential protein marker analysis was performed by western blot: melanosomes were validated using Tyrosinase, Dopachrome Tautomerase (DCT), and

Glycoprotein Non-Metastatic Melanoma Protein B (GPNMB), while small EV populations were characterized using CD63 and α-Enolase. Morphological examination employed both transmission electron microscopy (TEM) and standard electron microscopy (EM), and further particle tracking was achieved via fluorescence labeling and microscopy. Proteomic analyses of EVs utilized mass spectrometry, confirming the separation of exosomes and melanosomes, both of which were classified as EVs per MISEV2023 definitions. Based on this characterization, standardized functional comparison, and validated isolation procedure, all references to melanosomes in the current study designate them as large EVs, in alignment with contemporary guidelines and established methodological literature. Detailed procedures and functional characterization were performed as described in the referenced papers [28, 32, 33, 59], ensuring strict compliance with MISEV2023 protocol recommendations.

#### Cell culture

B16, and MNT-1 were all grown in complete DMEM (supplemented with 10% fetal bovine serum (FBS), 1 % penicillin/streptomycin, and 1 % L-glutamine, Gibco). Cells being used for melanosome purification were grown in vesicle-free FBS (FBS was centrifuged at 100,000rpm for 2 hours, only supernatant was added to culture media) and extra L-glutamine was added (to a final concentration of 2 %). All cells were grown at 37C with 5 % CO2. Primary human dermal fibroblasts were isolated as described previously [33].

#### ELISA

For the mouse serum ELISA, high-binding 96-well plates (Corning) were coated with 10 µg/mL of melanosomes or 5 µg/mL of purified protein and incubated overnight at 4 °C. Plates were then blocked with blocking buffer for 2 hours at room temperature. After blocking, the plates were washed with PBS containing 0.05% Tween-20 (PBST) and incubated with a 1:50 dilution of sera from melanosome-immunized mice for 1 hour at room temperature. For the mAbs binding to melanosomes ELISA, 96-well plates were coated with 10 µg/mL of melanosomes or with purified protein and incubated overnight at 4 °C. The plates were then washed with PBST and incubated with serial dilutions (as described in Figure legend) of mAbs, for 1 hour at room temperature. For the human serum ELISA, high-binding 96-well plates (Corning) were coated with 5 µg/mL of purified protein or 10 µg/mL of melanosomes and incubated overnight at 4 °C. Plates were then blocked with blocking buffer for 2 hours at room temperature and incubated with a 1:50 dilution of sera. Following washing with PBST, all plates were incubated with horseradish peroxidase (HRP)-conjugated anti-mouse IgG secondary antibody (0.16 µg/mL; Jackson ImmunoResearch) or anti-human IgG secondary antibody (0.16 µg/mL, Jackson ImmunoResearch) for 45 minutes at room temperature, followed by washing. TMB substrate (Abcam) was added, and optical density (OD) was measured at 650 nm. The Mouse IL-6 Uncoated ELISA Kit (Thermo Fisher, 88-7064-88) was performed according to the manufacturer’s instructions.

#### Flow Cytometry

Thawed spleen cells were resuspended in warm RPMI supplemented with 10% FBS, followed by centrifugation at 400 g for 5 minutes at 4 °C. The cell pellets were resuspended in FACS buffer and blocked with 1 µg/mL of anti-mouse CD16/32 (1:100, Thermo Fisher, 14-0161-85) for 15 minutes at 4 °C to prevent non-specific binding. Cells were then stained for 30 minutes at 4 °C with the following anti-mouse antibodies: anti-B220 PerCP-Cy5.5-conjugated (1:100, Biogems, 07131-70-100), anti-IgM BV421-conjugated (1:100, BioLegend, BLG-406506), anti-IgG1 BV421-conjugated (1:100, BioLegend, BLG-406616), anti-GL7 AF647-conjugated (1:100, BioLegend, BLG-144606), anti-FAS (CD95) PE-Vio770-conjugated (1:100, Milteny Biotec, 130-120-291), anti-CD4 BV421-conjugated (1:100, BioLegend, BLG-100438), anti-CD8a FITC-conjugated (1:100, BioLegend, BLG-100706), anti-NK1.1 PE-conjugated (1:100, Milteny Biotec, 130-120-511), anti-CD64 PE-Vio770-conjugated (1:50, Milteny Biotec, 130-119-659), anti-F4/80 APC-conjugated (1:50, Milteny Biotec, 130-116-525), anti-CD69 PE-Vio770-conjugated (1:10, Milteny Biotec, 130-103-977), anti-CD45 Brilliant Violet 711-conjugated (1:100, BioLegend, BLG-103147) antibodies, or anti-human/mouse antibodies: anti-HSP70 Alexa Fluor488-conjugated (1:100, BioLegend, BLG-648004), anti-GRP78 Alexa Fluor488-conjugated (1:100, Santa Cruz Biotechnology, SC-166490), anti-HSC70 Alexa Fluor488-conjugated (1:100, Santa Cruz Biotechnology, SC-7298). For HspA1A binding analysis by flow cytometry, Streptavidin Alexa Fluor 647 (BioLegend) was pre-incubated for 1 hour with biotinylated HspA1A. Subsequently, HspA1A (1 µg/mL per sample) was incubated with each monoclonal antibody (50 µg/mL) for 30 minutes before being added to 10^5^ B16 cells. Samples were analyzed using a CytoFLEX S4 flow cytometer (Beckman Coulter).

#### Single B cell sorting and sequencing

Murine B cells were isolated from the splenocytes of mice using mouse CD19 micro-beads (Miltenyi Biotec) according to the manufacturer’s instructions. B cells were mixed with anti-mouse CD16/CD32 (1:100, Thermo Fisher, 14-0161-85) and then stained with the following anti-mouse antibodies: anti-B220 PerCP-Cy5.5-conjugated (1:100, Biogems, 07131-70-100), anti-CD95 (FAS) PE-Vio770-conjugated (1:100, Milteny Biotec, 130-120-291), anti-GL7 AF647-conjugated (1:100, BioLegend, BLG-144606), IgM FITC-conjugated (1:100, BioLegend, BLG-406506) and IgG1 BV421-conjugated (1:100, BioLegend, BLG-406616). Single IgG1^+^ GC B cells were identified (B220^+^ CD95 (FAS)^+^ GL7^+^ IgM^−^ IgG1^+^) and were sorted using an Aria III sorter (Becton Dickinson) into a 96-well plate (4titude) containing 4 µl of lysis buffer. Following sorting, the plates were covered and immediately frozen on dry ice and transferred to −80 °C until further processing. Lysed cells were thawed on ice for RNA reverse transcription and PCR amplification as described previously [60–62]. Briefly, cDNA was synthesized using random hexamer primers (Invitrogen, 48190011), and SuperScript III Reverse Transcriptase (Invitrogen, 18080085). First round PCR products were used as a template for additional amplification by nested PCR with specific 5’ V and 3’ J primers containing restriction sites as previously described [61, 62]. The first round of PCR was carried out as: 98 °C for 30 seconds, 30 cycles of 98 °C for 30 seconds, 50 °C (Gamma) or 55 °C (Kappa) for 30 seconds, and 72 °C for 30 seconds. The second round of PCR was carried out under similar conditions, but the annealing temperature was changed to 57 °C (Gamma/Kappa) and the number of cycles reduced to 40 instead of 50. Second round PCR products were purified, sequenced, and annotated with IgBLAST [58].

#### Antibody sequence analysis

All PCR products were sequenced and analyzed for Ig gene usage, CDR3, and the number of V_H_/V_L_ somatic hyper mutations using IMGT (http://www.imgt.org) IgBLAST (http://www.ncbi.nlm.nih.gov/igblast/) databases. Expanded B cell clonal families were defined by ≥2 B cells exhibiting identical V_H_ J_H_ genes, and identical V_K_ J_K_ genes, displaying ≥75% homology in CDRH3, or in CDRK3 as previously reported [58].

#### Antibody and protein production

For mAb cloning, PCR products were purified (MACHEREY-NAGEL, 740609.250), digested with the appropriate restriction enzymes (New England Biolabs), and ligated (New England Biolabs, M0202L) into the corresponding expression vectors for IgG or IgK. Cloned mAb vectors for the IgG2a heavy chain and Kappa light chain were co-transfected into Expi293F cells at a ratio of 1:3 (H:K/L, Thermo Fisher Scientific Inc.) using the ExpiFectamine 293 Transfection Kit (Thermo Fisher Scientific Inc.). Seven days post-transfection, the cell supernatant was collected, filtered (0.22 μm), and incubated with protein A coated agarose beads (GE Life Sciences, 17519901) for 2 hours at room temperature. The beads were then loaded onto chromatography columns, washed, and eluted with 50 mM sodium phosphate (pH 3.0) into 1 M Tris-HCl (pH 8.0). Antibodies were buffer exchanged to PBS X1, aliquoted, and stored at −80 °C.

#### Live Imaging Experiments

B16, MNT1 melanoma cells or primary dermal fibroblasts (15 × 10³ cells) were seeded into a 96-well plate (Corning) with 200 µL of pre-warmed DMEM (Thermo Fisher Scientific), supplemented with 10% heat-inactivated FBS. For the direct effect assay, plates were pre-incubated for 30 minutes, followed by the addition of 25 µg/mL of each mAb directly to the cells. The plates were then immediately incubated and analyzed using the Incucyte® SX5 live-cell imaging system (Sartorius). For the external melanosome assay, 25 µg/mL of each mAb was incubated with 5 µg/mL of external melanosomes before being added to B16 cells. To measure apoptosis, Incucyte® Caspase-3/7 Green reagent (1:1000, Sartorius) was added to the culture medium, achieving a final concentration of 5 μM for the assay. Samples were monitored using the Incucyte® SX5 (Sartorius).

#### RNA extraction, library preparation, and sequencing

Total RNA was extracted and purified from B16 cells following a 24-hour treatment with mAbs: Mel321-31, Mel322-34, Mel321-35, and an isotype control (mGO.53). RNA extraction was performed using the PureLink™ RNA Mini Kit (Thermo Fisher Scientific) according to the manufacturer’s protocol. RNA quality and concentration were assessed using Qubit™ Fluorometer and Tapestation (Agilent Technologies). For transcriptome analysis, mRNA enrichment was performed using the NEBNext® Poly(A) mRNA Magnetic Isolation Module. Library preparation was carried out with the NEBNext® Ultra™ II Directional RNA Library Prep Kit, generating fragments of ∼ 350 bp (including adaptors). Sequencing was conducted at the Genomics Research Unit, supported by The Alfredo Federico Strauss Center for Computational Neuro-imaging, Faculty of Life Sciences, Tel Aviv University. Libraries were sequenced on the NextSeq2000 platform using a single-end 100 bp read strategy.

Raw reads were aligned to the mouse transcriptome and genome version GRCh39 with annotations from ENSEMBL release 106 using STAR aligner v.2.7.10a [63]. Counts per gene quantification was done using htseq-count v2.01 [64]. Genes with a sum of counts below 100 over all samples were filtered out. Gene expression was normalized per one million counts and log-transformed, and differential expression analysis was done with the PyDESeq2 package v 0.4.4 [65] with default parameters. Differentially expressed (DE) genes between the treatments were defined by applying a significance threshold of FDR corrected p-value<0.05.

Gene set enrichment analysis (GSEA) was performed using GSEA v.4.3.2 with the GSEA preranked tool. The Molecular Signature Database hallmark gene sets were used to perform pathway enrichment analysis.

#### Gene Ontology analysis

Differentially expressed genes were analyzed for their involvement in apoptosis-related pathways using Gene Ontology (GO) term annotations. The identified apoptosis-associated pathways include: GO:0006915, GO:0001783, GO:0042981, GO:0097192, GO:0043065, GO:0070059, GO:0006919, GO:0008630, GO:0008635, GO:0044346, GO:0070242, GO:1902262, GO:0097190, GO:1901030, GO:0008637, GO:0043524, GO:0043066, GO:0006919, GO:2001235, GO:0002904, GO:0044336, GO:0044337, GO:1902255, GO:2001243.

#### Western Blot analysis

Fresh melanosomes were lysed in radioimmunoprecipitation assay (RIPA) buffer (Thermo Fisher Scientific) and incubated on a rotator at 4°C for 1 hour. Lysates were centrifuged at 12,000 g for 15 minutes at 4°C, and protein concentration was determined using the Pierce™ BCA Protein Assay Kit (Thermo Fisher Scientific). 30 µg of each lysate was mixed with SDS reducing sample buffer, boiled for 5 minutes, separated by SDS-PAGE (Bio-Rad), and transferred onto a Trans-Blot Turbo™ nitrocellulose membrane (Bio-Rad). Membranes were blocked with 3% BSA in PBS (X1) for 1 hour at room temperature and incubated with specific primary antibodies (Mel321-31, Mel322-34, Mel321-35, and mGO.53) overnight at 4°C with gentle agitation. For HSP70 protein detection, 1 µg of each purified protein was mixed with SDS reducing sample buffer, boiled for 5 minutes, separated by SDS-PAGE (Bio-Rad), and transferred onto a Trans-Blot Turbo™ nitrocellulose membrane (Bio-Rad). Following transfer, membranes were blocked with 3% BSA in PBS (X1) for 1 hour at room temperature and incubated overnight at 4°C with gentle agitation with an anti-Avi tag monoclonal antibody (Avidity LLC). Detection was performed using an HRP-conjugated anti-mouse IgG secondary antibody (Jackson ImmunoResearch), Precision Protein™ StrepTactin-HRP Conjugate (1:5000, BioRad) and ECL reagent (Bio-Rad).

#### Immunoprecipitation and mass spectrometry

The samples were cleaved with trypsin and analyzed by LC-MSMS using the Q Exactive HF mass spectrometer, The data was analyzed with proteome Discoverer 2.4 software and the Sequest search engine against the specific database and a decoy database (in order to determine the false discovery rate, FDR). All the identified peptides were filtered with high confidence −1% FDR threshold. (*FDR = is the estimated fraction of false positives in a list of peptides). Quantitation was done by calculating the peak area of each peptide. The abundance of the protein is the sum of all associated peptide group abundances. T-test analysis between each group and the control was done using the Perseus software. P value<0.05. Supplementary file 1: Proteins with a logL fold change (FC) > 2 compared to the isotype control were highlighted in yellow in the Difference column. The Gene Name column includes color-coded annotations corresponding to different statistical tests applied. Differentially expressed proteins specific to the experimental groups, relative to the control (mGO.53, isotype control), are organized in separate tabs: one tab includes proteins consistently enriched across all three sample groups, while additional tabs categorize proteins specific to one or two experimental groups.

#### Amplification and cloning - HSP70

To produce HSP70, 0.1 × 10L B16 cells were seeded in a 24-well plate (Corning) and incubated for 24 hours. Total RNA was extracted using the PureLink™ RNA Mini Kit (Thermo Fisher Scientific) according to the manufacturer’s protocol. cDNA synthesis was performed using the qScript cDNA Synthesis Kit (Quanta Bio), and cDNA concentration was measured with Qubit™ dsDNA High-Sensitivity Assay Kit (Thermo Fisher Scientific). Amplification of HspA1A, HspA1L, HspA5, and HspA8 was carried out using the primers listed in Supplementary Table 2 and the KAPA HiFi HotStart ReadyMix (Roche). The PCR reaction consisted of 12.5 µL KAPA HiFi HotStart ReadyMix, 0.3 µM forward primer, 0.3 µM reverse primer, and 1 ng of template DNA, adjusted to 25 µL with DNase/RNase-free water. The thermocycling conditions were: 95 °C for 3 min (initial denaturation), 30 cycles of 98 °C for 20 s, 60 °C for 15 s, and 72 °C for 120 s, Final extension at 72 °C for 2 min. PCR products were purified (MACHEREY-NAGEL, 740609.250), digested with restriction enzymes (New England Biolabs), and ligated into the pcDNA 3.1 (+) mammalian expression vector (New England Biolabs, M0202L). Each construct included an N-terminal signal peptide (MKAPAVLAPGILVLLFTLVQRSNG) and two C-terminal tags: a hexa-histidine tag (His-tag, HHHHHHHH) and a site-specific biotinylation tag (AviTag, GLNDIFEAQKIEWHE). Phylogenetic trees were constructed based on the sequences using Geneious Prime 2024.0.

#### Transfection, production and protein isolation

Mammalian expression vectors containing the appropriate protein insert were transfected into Expi293F cells (Thermo Fisher Scientific) using the ExpiFectamine 293 Transfection Kit (Thermo Fisher Scientific). For antibody expression, a 1:3 heavy-to-light chain ratio was used. Seven days post-transfection, the cell supernatant was collected, filtered (0.22 μm), and incubated with Ni²L-NTA or Protein A-coated agarose beads (GE Life Sciences) for 2 hours at room temperature (RT). Proteins were eluted using 250 mM imidazole, buffer-exchanged into PBS (1×), aliquoted, and stored at −80°C. Antibodies were eluted using 50 mM sodium phosphate (pH 3.0) into 1 M Tris-HCl (pH 8.0) and underwent a similar purification process. When required, proteins were biotinylated using the BirA biotin-protein ligase kit (Avidity LLC, Colorado, USA) according to the manufacturer’s protocol.

#### Biacore (Surface Plasmon Resonance, SPR)

All Biacore experiments were performed at 25L°C with a Biacore T200 instrument. On a Series S Sensor Chip SA (GE Healthcare), 10 μg/mL CaptureSelect™ Biotin Anti-IgG-Fc (Multi-species) conjugate (Thermo Fisher Scientific) was immobilized at a flow rate of 10 μL/minute for 600 seconds. Over the chip-bound CaptureSelect, each mAb sample (Mel321-31, Mel322-34, Mel321-35, and isotype control mGO.53) at 1 μg/mL was injected at a flow rate of 10 μL/minute for 600 seconds at five sequential concentrations (31.25, 62.5, 125, 250, and 1000 nM). After each cycle, the chip was regenerated with 0.1 M glycine, pH 2, at a flow rate of 30 μL/minute for 90 seconds. Each antibody was tested in duplicate. Samples were diluted in HBS-EP buffer (10 mM HEPES, 150 mM NaCl, 3 mM EDTA, 0.05% Tween-20, pH 7.4). Sensorgrams were generated for each sample, and the curves were fitted to a 1:1 binding model in the BIAevaluation software using nonlinear regression. KD was calculated as the ratio of the dissociation (Kd) and association (Ka) rate constants, KD =LKd / Ka.

#### Alpha-Fold modeling

The protein complex and interactions formed between the mAbs (Mel321-31, Mel322-34, Mel321-35) and the proteins (HspA1A, HspA1L, HspA5 and HspA8) were modeled using the AlphaFold3 Server (https://alphafoldserver.com). Antibody-binding contact residues on HspA1A, HspA1L, HspA5, and HSPa8 were extracted using a Python [66] script (Supplementary file 1), applying a 5.0 Å distance cutoff. The extracted contact residues were then annotated using PyMOL 3.1. Model_0.cif was selected as the default structure. Only structures exhibited a confidence score (predicted TM-score) of pTM > 0.8 were used.

#### RNA isolation and Real-Time PCR

RNA was isolated using PureLink™ RNA (Thermo Fisher, 12183016) according to the manufacturer’s instructions. This was followed by cDNA synthesis using a qScript synthesis kit (Quanta bio), with the cDNA concentration assessed by NanoDrop. The qPCR mix was prepared using SYBR green (Quanta bio) with the primers described in Supplementary Table 3, Relative transcript expression was calculated using the ddCt method and all transcripts were normalized RPLPO.

#### Statistical analyses

All statistical analyses were performed using GraphPad Prism (version 9.5.1; GraphPad Software Inc., San Diego, CA, USA). Data were first assessed for normality using the Shapiro–Wilk test (for small sample sizes) or the D’Agostino–Pearson omnibus (K2) test (for larger datasets). For comparisons between two groups, we used: Unpaired two-tailed Student’s t-test for normally distributed data with equal variances. Unpaired two-tailed Welch’s t-test for normally distributed data with unequal variances. Mann–Whitney U test for non-normally distributed data. For comparisons among three or more groups, we used: One-way ANOVA, followed by Tukey’s multiple comparisons test for normally distributed data. Kruskal–Wallis test, followed by Dunn’s multiple comparisons test, for non-normally distributed data. For repeated measures data, we used: Repeated-measures one-way ANOVA, followed by Tukey’s multiple comparisons test. For Kaplan–Meier survival curves, we used the log-rank (Mantel–Cox) test to compare survival distributions between groups. For categorical data, we used: Fisher’s exact test for small sample sizes. All p-values were two-tailed, with p < 0.05 considered statistically significant. Adjustments for multiple comparisons were made using Tukey’s or Dunn’s post hoc tests, as appropriate. Data Presentation: Quantitative data are reported as mean ± standard deviation (s.d.) unless otherwise stated. Box plots display the median, interquartile range (IQR), and whiskers extending to 1.5× IQR. Individual data points are plotted where possible.

**Supplementary Figure Legends**

**Supplementary Figure 1. No differences in NK cells following melanosome-immunization. A.** Representative flow cytometry plot presents the gating strategy from a spleen of a melanosome-immunized mouse. Gating: lymphocytes → single cells and then immune cells were identified in Figure 1F-I. **B.** Flow cytometry plots of representative melanosome-immunized and PBS-immunized mice (Left), and corresponding frequencies (Right) of CD3^−^ NK1.1^+^ population in their spleens. Gating strategy is described in **A**; the experiment was repeated 2 times. Graphs present the meanL±Ls.d with each symbol represents an individual mouse. Statistical significance was determined using GraphPad Prism by unpaired two-sided Welch’s t-test. “ns” indicates non-statistically significant differences.

**Supplementary Figure 2. Sera responses against melanosomes are abolished following trypsinization.** Raw O.D. 650 nm values depicting the IgG response in melanosome-immunized mice (n=5) and PBS-immunized mice (n=5). Sera were collected 14 days post-immunization and assessed for IgG binding to melanosomes by ELISA. The graph shows raw O.D. 650 nm with each symbol represents an individual mouse. Statistical significance was determined using GraphPad Prism by one-way analysis of variance (ANOVA) with Tukey’s multiple comparisons post-test. ***, p < 0.001; ****, p < 0.0001; “ns” indicates non-statistically significant differences.

**Supplementary Figure 3. Elevated SHM rates and clonal expansion in GC B cells of immunized mice. A.** Number of SHMs (nucleotide changes) in V_H_ genes for each mouse clone (#Mel 18, #Mel 30, #Mel 31, and #Mel 32), derived from single sorted IgG1^+^ GC B cells on day 14 post-primary melanosome immunization. For reference, SHM data from V_H_ genes of single sorted GC B cells on day 14 post-OVA immunization is included on the right [39]. Graph presents the number with each symbol represents an individual sequence. **B.** Phylogenetic trees of the expanded clones shown in Figure 2E for melanosome-immunized mice (#18, #30, #31, #32). The legend for the trees is displayed at the bottom. Graph **A** presents the mean with each symbol represents an individual sequence. Statistical significance in **A** was determined using GraphPad Prism by one-way analysis of variance (ANOVA) with Tukey’s multiple comparisons post-test. **, p < 0.01; ****, p < 0.0001; “ns” indicates non-statistically significant differences.

**Supplementary Figure 4. TdT expression in B16 cells following melanosome addition and anti-melanosome mAb treatment.** Representative images of TdT-expressing B16 cells (red) under different treatment conditions: cells treated with isotype control, mGO.53 (25 µg/mL) plus 5 µg/mL of external melanosomes, Mel322-34 (25 µg/mL) plus 5 µg/mL of external melanosomes, and Mel321-35 (25 µg/mL) plus 5 µg/mL of external melanosomes. Scale bars: 400 µm.

**Supplementary Figure 5. Dose-dependent inhibition of B16 cell survival by anti-melanosome mAbs.** TdT-expressing B16 cells were incubated with Mel321-31, Mel322-34, or Mel321-35 at 50, 25, or 12.5 µg/mL. Cell fluorescence was measured after 48 hours. Data are presented as fold-change relative to cells treated with the isotype control antibody, mGO.53.

**Supplementary Figure 6. Immune profiling following treatment with the anti-melanosome mAb. A.** Flow cytometry plots showing the gating strategy used to identify CD3^−^ and CD3^+^ cells (Lymphocytes-singlets-CD3^−^/CD3^+^) in spleens of mAbs-treated mice. **B-D.** Left: Flow cytometry plots of immune populations in spleens of one representative Mel321-35 and isotype control treated mice. Right: Quantification of the gated population of Mel321-35 (n=9) and isotype control (n=10) treated mice. The immune populations include- **B:** Macrophages (CD3^−^ CD64^+^ F4/80^+^), **C:** NK cells (CD3^−^ NK1.1^+^), **D:** CD4^+^ T cells (CD3^+^ CD4^+^) and CD8^+^ T cells (CD3^+^ CD8^+^). **E.** Left: Flow cytometry plots of B220^+^ in TDLN of one representative Mel321-35 and isotype control treated mice. Right: Quantification of the gated populations of Mel321-35 (n=4) and isotype control (n=4) treated mice. **F.** Left: Flow cytometry plots of CD45^+^ in tumors of one representative Mel321-35 and isotype control treated mice. Right: Quantification of the gated populations of Mel321-35 (n=4) and isotype control (n=4) treated mice. Graphs **B-F** present the mean with each symbol represents an individual mouse. Statistical significance was determined using GraphPad Prism by unpaired two-sided Welch’s t-test in **B-F.** “ns” indicates non-statistically significant differences.

**Supplementary Figure 7. Mass spectrometry analysis following immunoprecipitation with anti-melanosome mAbs. A.** Mass spectrometry analysis of enriched melanosome proteins following immunoprecipitation with mAbs Mel321-31 (n=4), Mel322-34 (n=4), and Mel321-35 (n=4) compared to the isotype control, mGO.53 (n=4). Significant proteins (p < 0.05) with a positive fold change are color-coded: purple for Mel321-31, pink for Mel322-34, and red for Mel321-35. The y-axis represents the difference from the isotype control, mGO.53, and the x-axis represents the p-value. **B.** Western blot showing the purified HSP70 family members: HspA1A, HspA1L, HspA5 and HspA8 (0.5 µg each).

**Supplementary Figure 8. Affinity measurement of anti-melanosome mAbs to HspA1A.** SPR sensograms showing the binding interactions of the anti-melanosome mAbs Mel321-31, Mel322-34, and Mel321-35 (1 µg/mL) with immobilized HspA1A at sequential analyte concentrations of 31.25, 62.5, 125, 250, and 1000 nM.

**Supplementary Figure 9. Predicted epitopes of anti-melanosome mAbs.** Predicted binding epitopes of mAbs Mel321-31 (purple shades), Mel322-34 (pink shades), and Mel321-35 (red shades) to HspA1A, HspA1L, HspA5, and HspA8, as determined by AlphaFold3. The color scheme corresponds to each mAb as indicated. The substrate-binding domain (SBD) and nucleotide-binding domain (NBD) regions of HSP70 are marked. All predicted structures have a confidence score (predicted TM-score, pTM) > 0.8.

**Figure 10. Anti-HSP70 serum response correlates with better response to therapy.** Reactivity of IgG antibodies in sera from patients in the “Progressive Disease” group (red circles, n=19) and the “Complete Response” group (orange circles, n=19), assessed by ELISA against MNT-1, human melanosomes. Raw O.D.600 nm values were obtained after subtracting background binding to BSA. The dashed line represents the mean response of sera from healthy donors (n=4).

## Acknowledgments

This study is in the memory of Noga Kramer Yakov who passed away from melanoma aged 39. The study was funded by the Israel Science Foundation (ISF) grants [3136/22] and [638/23] to NTF; Binational Science Foundation (BSF) [01031771] to NTF; We thank Dr Sterna Germanis-Kaufman for support. NBS thanks Nilli and Dr Avi Shalev for their support. KPs research is supported in part by a fellowship from the Edmond J. Safra Center for Bioinformatics at Tel-Aviv University. YW is supported by a Melanoma Research Alliance grant (no.937368), the Rosetrees Trust (no. MYIA\100002), a research grant from Pfizer, and the Lemelbaum family. CL acknowledge that this work was Funded by the European Research Council (ERC) under the European Union’s Horizon 2020 research and innovation program European Union. Views and opinions expressed are however those of the author(s) only and do not necessarily reflect those of the European Union or the ERCEA. Neither the European Union nor the granting authority can be held responsible for them (grant agreement no. 726225). We thank Roi Balaban for helping with the in vivo experiments. We thank Sebastien Apcher, Institute Gustave Roussy, France, for his conceptual contribution to the idea of immunizing mice with melanosomes and for early discussions with Carmit Levy that led to the initiation of the current study. Figures: 1A, 2A, 4A were created using BioRender.com.

## Authors’ contribution

NBS designed and conducted the experiments, analyzed data, prepared figures, and co-wrote the manuscript with NTF. SP performed mice melanosome immunizations and tumor challenge, that were repeated by NBS. LA assisted NBS with in vivo experiments. RY assisted NBS with ELISA and Flow Cytometry experiments. PM, RP and SM contributed to melanosome production and cell line culturing. DFC and TH contributed to proteomic analysis of the mAb immunoprecipitation experiments. OM and MC assisted NBS with mRNA-seq data analysis. KP contributed to the statistical analysis. YW and RSF recruited the melanoma patients, provided the human sera samples and applied for the IRB approvals. SA contributed conceptually to the idea of immunizing mice with melanosomes. CL together with NBS and NTF participated in the initiation, ideation and conceptualization of the study. NTF applied for TAU ethic approvals, conceived and supervised the study, analyzed data, prepared figures, and co-wrote the manuscript with NBS.

## Inclusion and Ethics statement

All the authors meet the authorship criteria and gave their consent to be listed as authors on this manuscript. We are committed to fostering an inclusive and diverse research environment. Our collaborative team comprises researchers from different cultural, gender, and academic backgrounds, and we actively promote equity in scientific collaboration and mentorship. Our research adheres to the principles of diversity, equity, and inclusion, and we strive to provide a welcoming and supportive space for all individuals.

