## Supplementary Tables for "Decoy Antibodies Block Extracellular HSP70, Prevent Self Signaling and Inhibit Melanoma Cell Survival"

**Supplementary Table 1** – Mouse primers - single cells sorting - *In vivo*

| HP1F-mLEADER-1 | GTAACACTTTTAAATGGTATCCAGTGT |
| --- | --- |
| HP1F-mLEADER-2 | GTCCTAATTTTAAAAGGTGTCCAGTGT |
| HP1F-mLEADER-3 | AGCAACAGCTACAGGTGTCCACTCC |
| HP1F-mLEADER-4 | TCTTCTTCCTGTCAGTAACTRCAGG |
| HP1F-mLEADER-5 | TTGCTATTCCTGATGGCAGCTGCCCAA |
| HP1F-mLEADER-6 | GTGACATTCCCAAGCTGTGTCCTRTCC |
| HP1F-mLEADER-7 | TGTACCTGTTGACAGTCGTTCCTGG |
| HP1F-mLEADER-8 | GTTTTTTATCAAGGTGTGCATTGT |
| HP1F-mLEADER-9 | CTATTCCTGATGGCAGCTGCCCAAAG |
| IgG HP1R-3′-Cγ1-outer | GGAAGGTGTGCACACCGCTGGAC |
| HP2F-5′-MsVHE | GGGAATTCGAGGTGCAGCTGCAGGAGTCTGG |
| IgG HP2R-3′-Cγ1-inner | GCTCAGGGAAATAGCCCTTGAC |

**Supplementary Table 2** – HSP70 primers- Cloning

| HSPA1A-Fw | CCTCTAGACTCGAGCGGCCGCATGAAGGCCCCCGCTGTGCTTGCACCTGGCATCCTCGTGCTCCTGTTTACCTTGGTGCAGAGGAGCAATGGGATGGCCAAGAACACGGCGAT |
| --- | --- |
| HSPA1A-Rev | CAGCGGTTTAAACTTAAGCTTTCATTCGTGCCACTCAATTTTTTGCGCTTCAAAGATGTCGTTCAGGCCGTGGTGGTGGTGGTGGTGGTGGTGATCCACCTCCTCGATGGTGG |
| HSPA1L-Fw | CCTCTAGACTCGAGCGGCCGCATGAAGGCCCCCGCTGTGCTTGCACCTGGCATCCTCGTGCTCCTGTTTACCTTGGTGCAGAGGAGCAATGGGATGGCTGCTAATAAAGGAATGGCG |
| HSPA1L-Rev | CAGCGGTTTAAACTTAAGCTTTCATTCGTGCCACTCAATTTTTTGCGCTTCAAAGATGTCGTTCAGGCCGTGGTGGTGGTGGTGGTGGTGGTGATCTACTTCCTCGATGGTAGGGC |
| HSPA5-Fw | CCTCTAGACTCGAGCGGCCGCATGAAGGCCCCCGCTGTGCTTGCACCTGGCATCCTCGTGCTCCTGTTTACCTTGGTGCAGAGGAGCAATGGGGGATGATGAAGTTCACTGTGGTGGC |
| HSPA5-Rev | CAGCGGTTTAAACTTAAGCTTTCATTCGTGCCACTCAATTTTTTGCGCTTCAAAGATGTCGTTCAGGCCGTGGTGGTGGTGGTGGTGGTGGTGCAACTCATCTTTTTCTGATGTATCCTCTTCACC |
| HSPA8-Fw | CCTCTAGACTCGAGCGGCCGCATGAAGGCCCCCGCTGTGCTTGCACCTGGCATCCTCGTGCTCCTGTTTACCTTGGTGCAGAGGAGCAATGGGATGTCTAAGGGACCTGCAGTTGG |
| HSPA8-Rev | CAGCGGTTTAAACTTAAGCTTTCATTCGTGCCACTCAATTTTTTGCGCTTCAAAGATGTCGTTCAGGCCGTGGTGGTGGTGGTGGTGGTGGTGATCCACCTCTTCAATGGTGGGG |

**Supplementary Table 3** – Real-Time PCR primers

| Rplp0_human_Fw | TGGTCTATCCAGCAGGTTTCGA |
| --- | --- |
| Rplp0_human_Re | ACAGACACTCGCACACTTCCGG |
| Hu_A1A_Fw | ACCTTCGACGTGTCCATCCTGA |
| Hu_A1A_Rev | TCCTCCACGAAGTGGTTCACCA |
| Hu_A1L_Fw | CAAACTGAAGGGAAGGCGTC |
| Hu_A1L_Rev | TCGATGCCTATGGCGATTCC |
| Hu_A5_Fw | CTGTCCAGGCTGGTGTGCTCT |
| Hu_A5_Rev | CTTGGTAGGCACCACTGTGTTC |
| Hu_A8_Fw | TTATTGGAGCCAGGCCTACAC |
| Hu_A8_Rev | GTGCTGGAAAACACCCACAC |
